# *Vitis vinifera* plants edited in *DMR6* genes show improved resistance to downy mildew

**DOI:** 10.1101/2022.04.19.488768

**Authors:** Lisa Giacomelli, Tieme Zeilmaker, Simone Scintilla, Umberto Salvagnin, Jeroen Rouppe van der Voort, Claudio Moser

## Abstract

The production and cultivation of vines (*Vitis vinifera*) tolerant or resistant to diseases such as downy mildew (DM) is a promising strategy to reduce fungicides and help viticulture sustainability. In many crops, generation of knock-out mutants in host genes controlling susceptibility to DM, such as *Downy Mildew Resistant 6* (*DMR6*) is a strategy of proven success to obtain resistant plants, while the effect of mutations in *DMR6* genes has yet to be demonstrated in grapevine. In addition, small mutations in genes governing important traits can be obtained by gene-editing while maintaining the genetic background of commercially important clones. Moreover, very recent advances in the technology of gene-editing allowed to produce non-transgenic grapevine mutants, by regeneration of protoplasts previously edited with the CRISPR-Cas9 ribonucleoprotein. This approach may revolutionize the production of new grapevine varieties and clones, but it requires knowledge on the targets, and an extensive evaluation of the impact of their mutation on plant phenotype and fitness. In this work we generated single and double knock-out mutants in *DMR6* susceptibility (S) genes in multiple grapevine cultivars with improved resistance to DM.

## INTRODUCTION

Cultivated grapevine (*Vitis vinifera* L.) is highly susceptible to several diseases (Vezzulli et al., 2022), typically caused by fungi and oomycetes. In humid weather, the most devastating disease is downy mildew (DM)–caused by the oomycete *Plasmopara viticola*, whereas in hot and dry climates powdery mildew (PM) *i*s mostly an issue–caused by the ascomycete *Erysiphe necator*. Through lengthy breeding programs, resistant cultivars and hybrids with good oenological attitudes were obtained mainly by introgression of resistance (R) genes. This allowed to reduce the yearly applications of fungicides, a condition that turned favorable to the emergence of new diseases: among them is black rot, caused by the ascomycete *Guignardia bidwellii* (Töpfer et al., 2011). Therefore, an effective reduction of chemical treatments in the vineyard can be in principle achieved with vines that are resistant to multiple pathogens. Susceptibility (S) factors are encoded by plant genes, and are required for the establishment of a particular disease (van Schie and Takken, 2014). S genes facilitate infection and support compatible interaction with a pathogen, are generally recessive, and their mutations generally produce resistance towards more than one pathogen. This is the case for mutations in the *Downy mildew resistant 6* (*DMR6*) gene, which produces resistance to oomycetes as well as fungi and bacteria in several crops, including tomato (de Toledo Thomazella et al., 2021), potato (Kieu et al., 2021), banana (Tripathi et al., 2021), apple (Shan et al., 2021), sweet basil (Hasley et al., 2021), and citrus (Parajuli et al., 2022). Its role as an S gene has been well established in *Arabidopsis thaliana*: the *dmr6* mutant displays resistance to the DM-causative agent *Hyaloperonospora arabidopsidis*, to the oomycete *Phytophthora capsici*, and the bacterium *Pseudomonas syringae*. (Zeilmaker et al., 2015)

*AtDMR6* encodes a 2-oxoglutarate (2OG)-Fe(II) oxygenase, which functions as a negative regulator of immunity. In Arabidopsis, two additional DMR6-Like Oxygenases (AtDLO1 and AtDLO2) restore DM susceptibility when over-expressed in the *dmr6* mutant (Van Damme et al., 2008; Zeilmaker et al., 2015). Functionally, AtDMR6 and AtDLO1 are respectively a salicylic acid (SA) 5-hydroxylase and a SA 3-hydroxylase (Zhang et al., 2013; Zhang et al., 2017), with an effect on SA homeostasis and senescence.

In grapevine, the DMR6-DLO family consists of two highly similar genes (*VviDMR6-1* and *VviDMR6-2*) and three encoding DMR6-like oxygenases (*VviDLO1, 2* and *3*). A recent analysis of the grapevine *DMR6* and *DLO* networks of transcriptionally co-regulated genes revealed a consistent group of defense-associated genes that are co-regulated with *VviDMR6-1* and *VviDMR6-2*, and that over-expression of *VviDMR6-1* restores susceptibility to DM of the of the Arabidopsis *dmr6-1* resistant mutant (Pirrello et al., 2022).

In this work, we generated single and double mutants in *VviDMR6-1* and *VviDMR6-2* by gene editing using CRISPR/Cas9. We show that mutations in *VviDMR6-1* and *VviDMR6-2* reduce susceptibility to DM in Crimson seedless.

## MATERIALS AND METHODS

### CRISPR/Cas9 construct

Guide RNAs to specifically target grapevine *DMR6* genes were designed on the Pinot PN4004 reference genome (Jaillon et al., 2007) using the CRISPR-P software tool (Lei et al., 2014). The selected sgRNAs were cloned using the Golden Gate cloning method into an expression vector behind the U6 promoter, and subsequently into a binary vector containing a domesticated Cas9 driven by a double 35S promoter, and obtained from the Addgene plasmid repository (www.addgene.org). Target and PAM site regions were sequenced by Sanger and Next Generation Sequencing (NGS) in the different cultivars to check for absence of polymorphisms. The sgRNAs used in this work are based on the following target sites: GCCGATGCTTGCAGGCTCTA (DM1a) and GTCCTTGCCGAGGTCGATTA (DM1b) for the *VviDMR6-1* gene (VIT_16s0098g00860); GGGCTCGATCGTCACAACTC (DM2a), GATGTAGTTCTCCGGCAAAG (DM2b), and GGAGGATTGGAGGGCCACTC (DM2c) for the *VviDMR6-2* gene (VIT_13s0047g00210). Guides DM1a and DM1b were used respectively in the constructs pDM1a and pDM1b to edit *VviDMR6-1* alone; guides DM2a and DM2b were used in the constructs pDM2a and pDM2b respectively to edit *VviDMR6*-2 alone. In addition, vectors for simultaneous expression of two sgRNAs were constructed: pDM1a2a (with guides DM1a and DM2a), pDM1a2b (with guides DM1a and DM2b), and pDM1a2c (with guides DM1a and DM2c) to edit simultaneously *VviDMR6-1* and *VviDMR6-2* (table 1).

**Table 1:** Transgenic grapevine plants regenerated from transformed calli and sequenced.

| Plants sequenced and edited in *VviDMR6-1* sgRNA1 (target DM1A) |  |  |  |  |  |  |  |
| --- | --- | --- | --- | --- | --- | --- | --- |
| Plasmid | Cultivar | Sequenced | WT | Edited | Edited >99% | Edited 90-99% | Edited 1-90% |
| pDM1a2a | Chardonnay | 23 | 15 | 8 | 0 | 0 | 8 |
| pDM1a2a | Sugraone | 169 | 25 | 144 | 5 | 2 | 137 |
| pDM1a | Crimson S. | 71 | 28 | 43 | 3 | 1 | 39 |
| pDM1a | Microvine | 1 | 0 | 1 | 0 | 0 | 1 |
| pDM1a | Sugraone | 164 | 55 | 109 | 30 | 8 | 71 |
| pDM1a2c | Crimson S. | 87 | 25 | 62 | 5 | 0 | 57 |
| pDM1a2c | Sugraone | 26 | 9 | 17 | 5 | 0 | 12 |
| Total |  | 541 | 157 | 384 | 48 | 11 | 325 |
| Plants sequenced and edited at *VviDMR6-1* sgRNA2 (target DM1b) |  |  |  |  |  |  |  |
| Plasmid | Cultivar | Sequenced | WT | Edited | Edited >99% | Edited 90-99% | Edited 1-90% |
| pDM1b | Sugraone | 100 | 9 | 91 | 4 | 8 | 79 |
| Total |  | 100 | 9 | 91 | 4 | 8 | 79 |
| Plants sequenced and edited in *VviDMR6-2* sgRNA1 (target DM2a) |  |  |  |  |  |  |  |
| Plasmid | Cultivar | Sequenced | WT | Edited | Edited >99% | Edited 90-99% | Edited 1-90% |
| pDM1a2a | Chardonnay | 21 | 21 | 0 | 0 | 0 | 0 |
| pDM1a2a | Sugraone | 143 | 143 | 0 | 0 | 0 | 0 |
| pDM2a | Sugraone | 2 | 1 | 1 | 0 | 0 | 1 |
| pDM2a | Crimson S. | 92 | 81 | 11 | 0 | 0 | 11 |
| pDM2a | Merlot | 1 | 0 | 1 | 0 | 0 | 1 |
| pDM2a | Thompson S. | 52 | 11 | 41 | 1 | 0 | 40 |
| Total |  | 311 | 257 | 54 | 1 | 0 | 53 |
| Plants sequenced and edited in *VviDMR6-2* sgRNA2 (target DM2b) |  |  |  |  |  |  |  |
| Plasmid | Cultivar | Sequenced | WT | Edited | Edited >99% | Edited 90-99% | Edited 1-90% |
| pDM2b | Crimson S. | 65 | 63 | 2 | 0 | 0 | 2 |
| pDM2b | Sugraone | 133 | 128 | 5 | 0 | 0 | 5 |
| Total |  | 198 | 191 | 7 | 0 | 0 | 7 |
| Plants sequenced and edited in *VviDMR6-2* sgRNA3 (target DM2c) |  |  |  |  |  |  |  |
| Plasmid | Cultivar | Sequenced | WT | Edited | Edited >99% | Edited 90-99% | Edited 1-90% |
| pDM1a2c | Crimson S. | 92 | 4 | 88 | 78 | 2 | 8 |
| pDM1a2c | Sugraone | 29 | 1 | 28 | 25 | 0 | 3 |
| Total |  | 121 | 5 | 116 | 103 | 2 | 11 |

### Plant material and gene transfer

Embryogenic calli were obtained from cultures of ovaries and anthers of *V. vinifera* cv. Sugraone, Crimson seedless, Thompson seedless, and microvine 04C023V0006 (Chaïb et al., 2010) according to Martinelli et al., 2001. These calli were cultivated in absence of selection to regenerate the wild type plants used in this work, and in parallel they were transformed by co-cultivation with *Agrobacterium tumefaciens* EHA105 carrying the proper binary vector, essentially as described by Dalla Costa et al., 2016. Embryogenesis was induced in the dark on NN solid medium (Nitsch and Nitsch, 1969) supplemented with 1 g/L activated charcoal, 45 g/L sucrose, 150 μg/L kanamycin, 1 mg/L timentin, 0.9 μM 6-benzylaminopurine, 11.4 μM indole 3-acetic acid, and 10 μM beta-naphtoxyacetic acid. Depending on the cultivar, embryos were cultivated on NN solid medium, 15 g/L sucrose, and 25 μg/L kanamycin in the light (16 h photoperiod) with or without the supplement of growth-hormones (4.5 μM 6-benzylaminopurine, 5 μM 3-Indolebutyric acid). Transgenic lines were screened by PCR with NPTii specific primers (GCCAACGCTATGTCCTGATA, and ACAATCGGCTGCTCTGATG). Edited lines were sequenced on a MiSeq system (Illumina, San Diego, CA, USA) and reads were analyzed through the CRISPResso platform (Pinello et al., 2016) using standard settings for filtering of low-quality reads and trimming of adapter sequences.

In addition, the transgene-free *dmr6-2* mutant H1D was obtained via single-cell technology by editing of protoplasts of cv. Crimson seedless with the ribonucleoprotein complex: TrueCut Cas9 protein (Thermo Fisher Scientific, Waltham, MA, USA) and sgRNA DM2c, as described in Scintilla et al., 2021.

### Plant growth

Regenerated plants (both wild type and edited) were propagated on NN medium (Nitsch and Nitsch, 1969) with 15 g/L sucrose and maintained in a climate-controlled chamber (16 h light photoperiod, 23 °C, 60% relative humidity). Plants were acclimatized in rooting soil with low percentage of pumice, and grown in a clean environment under LED light (400 μmol photons m^-2^ S^-1^ at the plant-tip height, 16 h light photoperiod).

### DM assay

*P. viticola* (Berk. & M.A. Curtis) Berl. & De Toni was propagated on susceptible plants in the greenhouse. Symptomatic plants were placed in the dark at 100% relative humidity (RH) overnight to induce sporulation. The inoculum was prepared by collecting sporangia from leaves in cold water to a concentration of 10^5 sporangia/mL. The DM assay was performed by inoculation of leaf discs harvested actively growing plants (untreated and healthy) with 15-20 leaves per stem. Leaf discs (0.8 cm diameter) were cut from the third to ninth leaf of a growing stem with a cork borer and laid–abaxial-side up–on four sheets of wet absorbing-paper in Microbox containers (Sac O2, Nevele, Belgium) in the light at 25 °C. To account for ontogenic resistance, leaves were grouped according to their age, as determined by their position on the stem. Leaves of the same age group were treated together and compared within the group. Severity, (calculated as percentage of the leaf-disc area showing sporulation) and incidence were evaluated at 7 dpi with the ImageJ software. The average severity of the wild type was used as internal control for each experiment and leaf age, and used as normalizer to compare data collected from multiple experiments. Statistically significant differences were evaluated in Python by ANOVA Tukey-HSD in Python within the Google Colab platform.

## RESULTS AND DISCUSSION

### Generation of mutants

Kanamycin-resistant mutants of seven cultivars were selected (table 1), and 994 plants were sequenced: 236 plants transformed with pDM1a, 100 with pDM1b, 147 with pDM2a, 198 with pDM2b, 192 with pDM1a2a, and 121 with pMD1a2c. Due to the recessive nature of the resistance phenotype, we selected completely edited plants preferably with mutations causing a frameshift of the reading frame and resulting in an early stop codon. Transformed plants were discriminated by NGS in two groups: those with editing in at least 90% of the reads (blue and green sectors in pies of figure 1) were further selected and propagated, whereas plants with editing in less than 90% of the reads were discarded. The great majority of transformed plants fall in this second group and include transformed but non-edited plants, heterozygous plants, as well as edited chimeras with portions of non-edited sequences. The stability of mutations was checked by sequencing of cuttings over time. In general, we observed that plants retaining less than 1% of non-edited DNA–comparable to sequencing noise–generated fully edited plants also after multiple cycles of propagations via cuttings. This is the case of lines 51, B95, and the double mutant O79 (figure S1). Differently, plants 58 and B18 retaining 1-10% of non-edited DNA showed an unstable genotype in propagated cuttings (figure S1). Therefore, to ensure genetic stability of cuttings and ensure complete gene knock-out, we further selected plants that were edited in more than 99% of the reads, and refer to these lines as completely edited (blue sectors of pies in figure 1). A selection of these plants was maintained and further analyzed.

**Figure 1:**
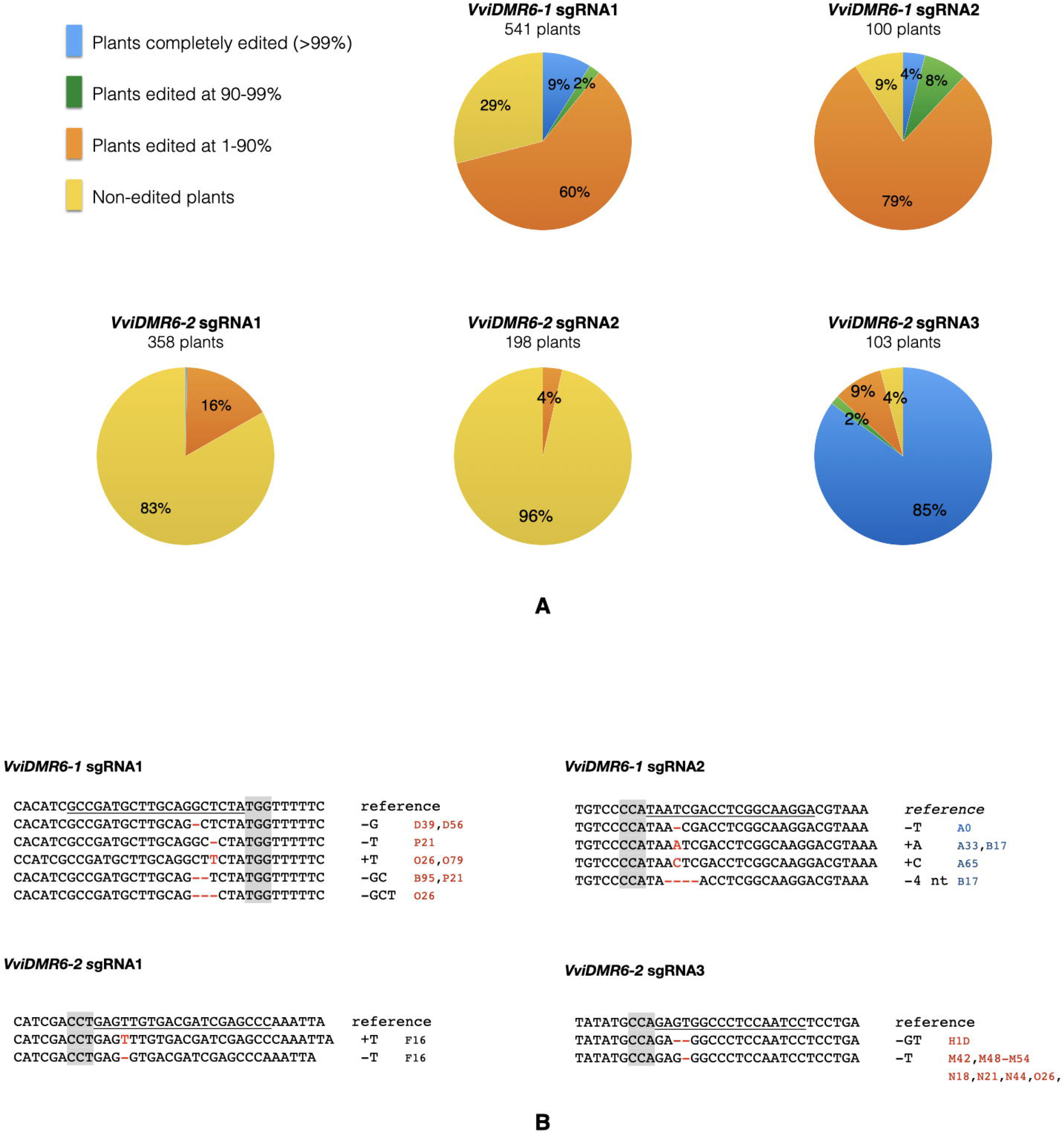
Genotyped edited plants. (A) Grouping of edited plants based on the fraction of edited reads in NGS sequencing. Completely edited plants were defined as those with more of 99% of edited reads. (B) Editing types of completely edited plants selected for phenotypic analysis. The reference sequence refers to that of cv. Sugraone and Crimson seedless, which are identical in these target regions. Mutations are indicated in red, and a list of lines showing the reported mutations is provided. Lines of cv. Sugraone are indicated in blue, and lines of cv. Crimson seedless are indicated in red. The genetic background of line F16 is cv. Thompson seedless.

### Chimerism

Genetically modified grapevines have been so far mostly produced by gene-transfer of embryogenic callus, since the technology to generate transgene-free grapevines from edited protoplasts is very recent (Scintilla et al., 2021). In *Agrobacterium*-mediated transformation of embryogenic callus, the formation of genetic chimeras is rather common (Nakajima et al., 2017) since it is based on cocultivation of plant cells and the bacterium. While the selection against *Agrobacterium* commences, a mixture of undifferentiated grapevine cells and pro-embryonic clusters are induced to differentiate and divide. A chimera is generated when gene-transfer (and Cas9-mediated mutation) occurs in cells that are already part of a pro-embryonic cluster. In the case of CRISPR/Cas9 editing, an additional level of complexity can contribute to the generation of chimeras, which depends on the cutting efficiency the CRISPR/Cas9 machinery, and on the outcome of the endogenous repairing process. In this work, we relied on the production of a large number of regenerated plants to select completely-edited, non-chimeric grapevine mutants in one or even two genes simultaneously. In this regard, we counted on NGS to detect chimeras and small portions of non-edited sequenced, which would likely go undetected by Sanger sequencing. Detection of even traces of wild type sectors in transgenic plants is especially important when they may mask the phenotype of a recessive mutation, such as in S genes.

### Editing efficiency

The editing efficiency of a specific sgRNA is known to depend on its sequence and secondary structure, as amply discussed in the literature (Liang et al., 2016; Graf et al., 2019; Uniyal et al., 2019). In our case, the influence of the sgRNA on the editing efficiency is evident for the *VviDMR6-2* gene, where sgRNAs with targets DM2a and DM2b were virtually ineffective, whereas the sgRNA based on target DM2c was extremely efficient: 85% of the screened plants were completely edited (figure 1), and mostly monoallelic. Given its high efficiency, this guide was also used by Scintilla and collaborators to produce non-transgenic *dmr6-2* mutants (line H1D) from protoplasts via the single-cell technology (Scintilla et al., 2021).

### dmr6 mutants are less susceptible to DM than wild-type

Young potted plants of selected edited genotypes (*dmr6-1, dmr6-2* and *dmr6-1_dmr6-2*) did not show any evident growth phenotype in their juvenile stage in controlled conditions (figure **2**). Leaf discs of these plants were artificially inoculated with *P. viticola*, and the severity of sporulation was scored at 7 dpi.

**Figure 2:**
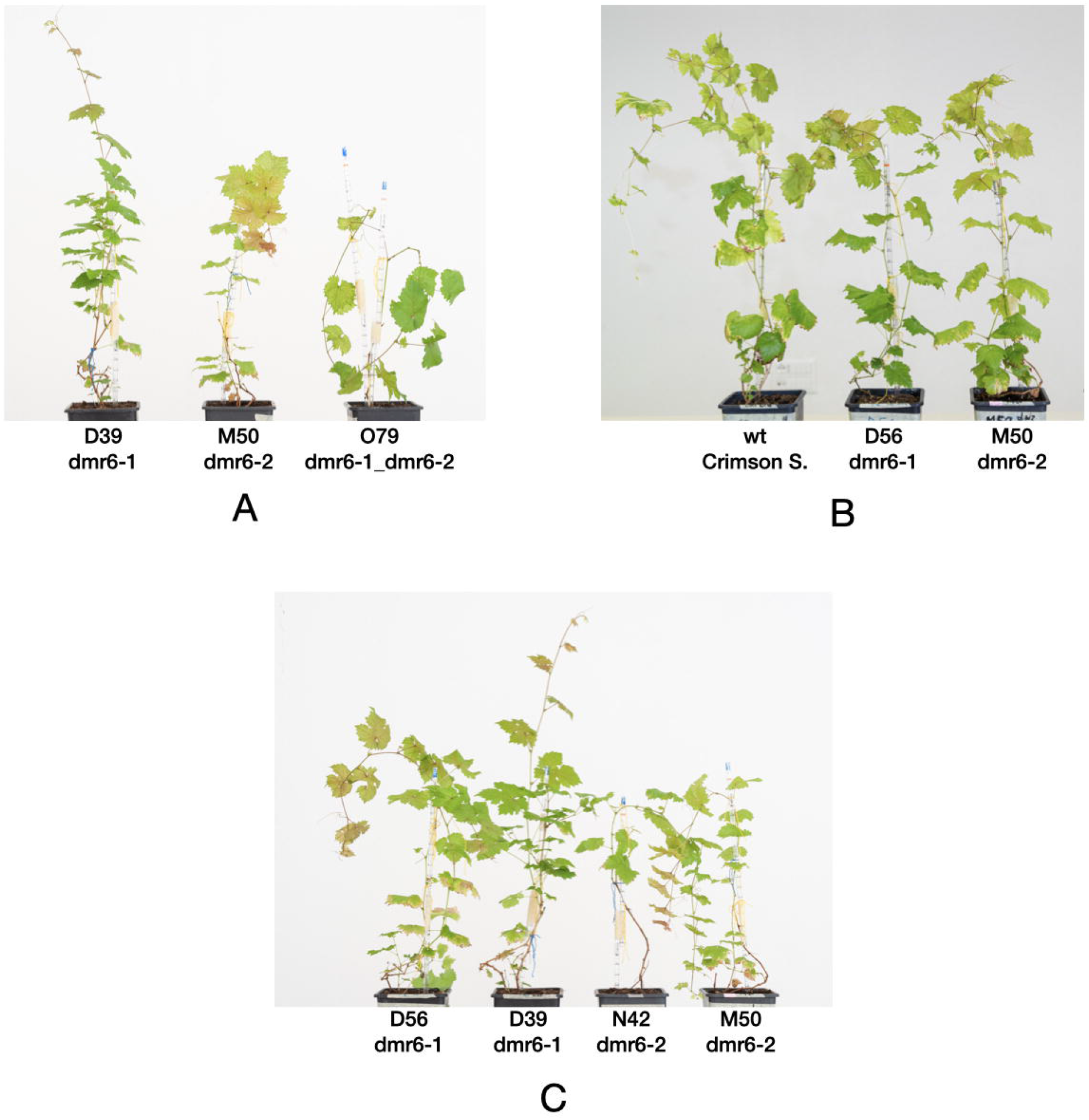
Pictures of edited mutants. (A) Comparison between lines of single and double mutants. (B) Comparison between wild-type and lines of single mutants. (C) Comparison between independent transgenic lines of single *dmr6-1* mutants (lines D39 and D56), and single *dmr6-2* mutants (lines N42 and M50). Pictures were taken about four weeks after acclimatization of plants to soil.

Single *dmr6-1* and *dmr6-2* mutants as well as *dmr6-1_dmr6-2* double mutants showed reduced relative severity as compared to the wild type (figure 3) in cv. Crimson seedless, and the difference between mutant genotypes and the wild type is significant. The analyzed plants were: *dmr6-1* line D39, *dmr6-2* lines M54, and H1D, and double mutants M42, O26 and O79. All lines are transgenics with the exception of the *dmr6-2* mutant H1D, which was produced by the single-cell technology (Scintilla et al., 2021) consisting in editing of protoplasts with the CRISPR/Cas9 ribonucleoprotein, followed by regeneration of non-transgenic edited plants. No significant difference in susceptibility was observed between independent lines within the same genotype group (figure 3).

**Figure 3:**
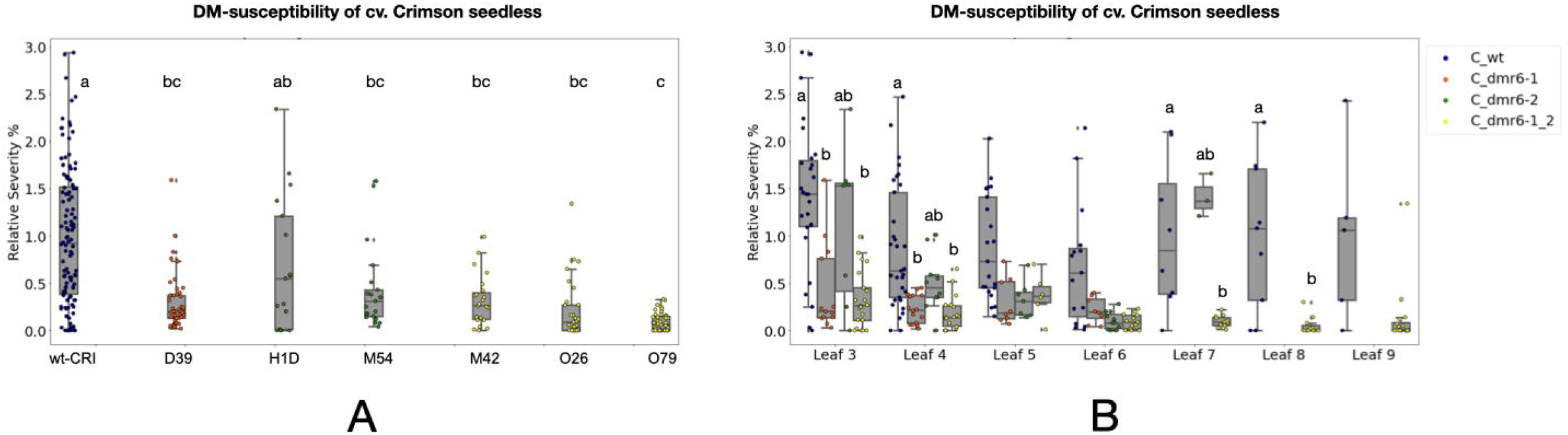
Relative severity of mutants on detached leaf discs. The relative severity (%) is plotted per genotype (A), and per leaf age (B). Leaf were numbered starting from the first flat leaf form the top of the shoot. Statistically significant differences according to Anova Tukey HSD (p>0.05) are indicated by different letters. The genotypes are indicated by color: the wild type in blue, the single *dmr6-1* mutant in red, the single *dmr6-2* mutant in green, and the double mutants in yellow.

## CONCLUSIONS

This study shows that grapevine *DMR6* genes are promising candidates for gene-editing to obtain DM-resistant vines, although further experiments are needed to confirm their potential as S genes, together with a thorough evaluation of the phenotype of these mutants. In combination with the recent technology that allows to produce edited plants without any trace of foreign DNA, this opens up new roads towards a more sustainable viticulture.

## ACKNOWLEDGMENTS

We thank Dr. Marco Moretto for help with the Python Colab platform, and Dr. Oscar Giovannini for maintenance of the *P. viticola* inoculum.

---

**Figure S1: Sequencing of propagated cuttings.**

The figure shows the CRISPResso output of selected edited plants and their cuttings propagated over time. A red arrow indicates the non-edited DNA retained by the mother plants and their cuttings.

