## Supplementary material for "*Vitis vinifera* plants edited in *DMR6* genes show improved resistance to downy mildew": Figure S1

**Figure S1: Sequencing of propagated cuttings.**

The figure shows the CRISPResso output of selected edited plants and their cuttings propagated over time. A red arrow indicates the non-edited DNA retained by the mother plants and their cuttings.

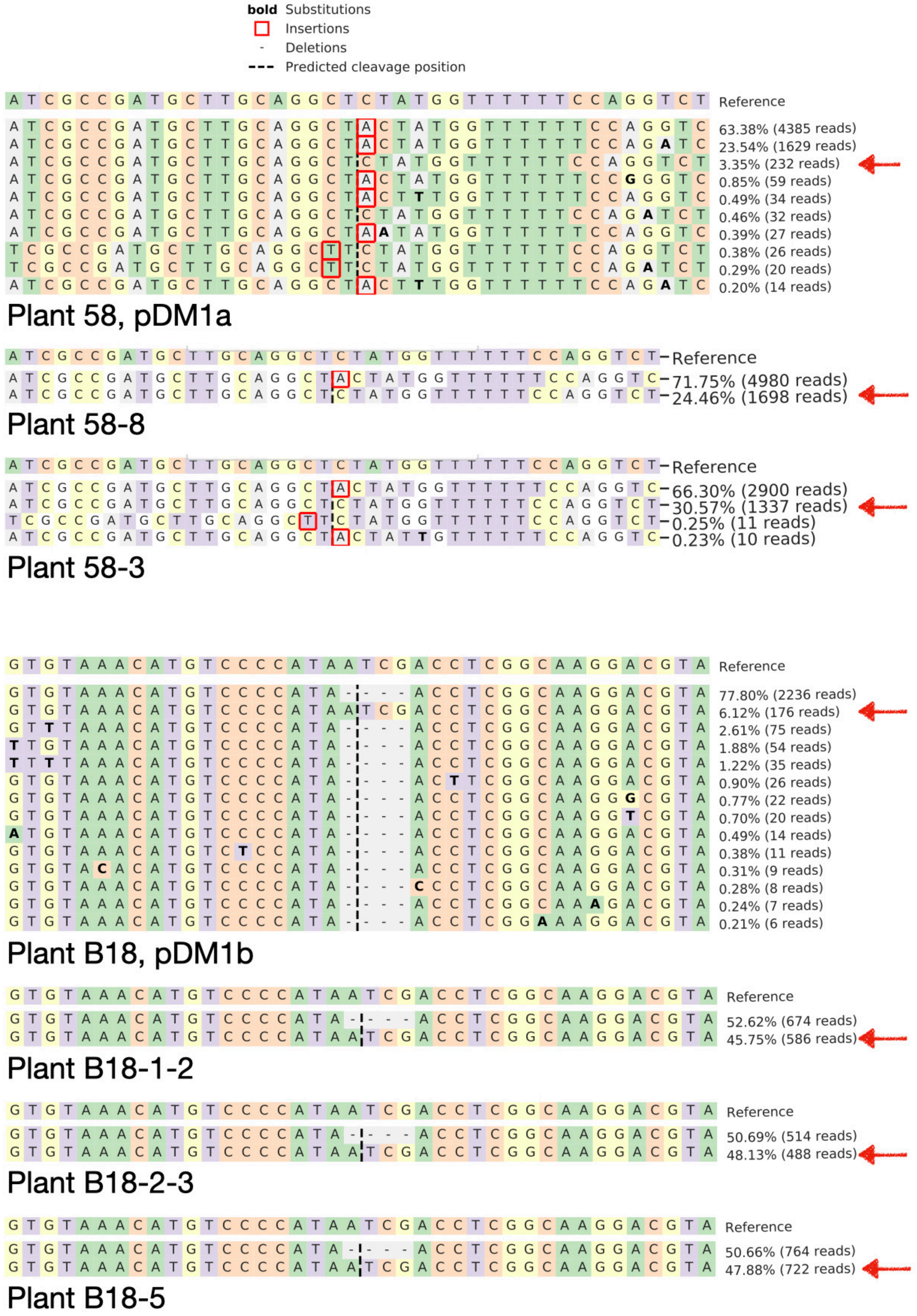

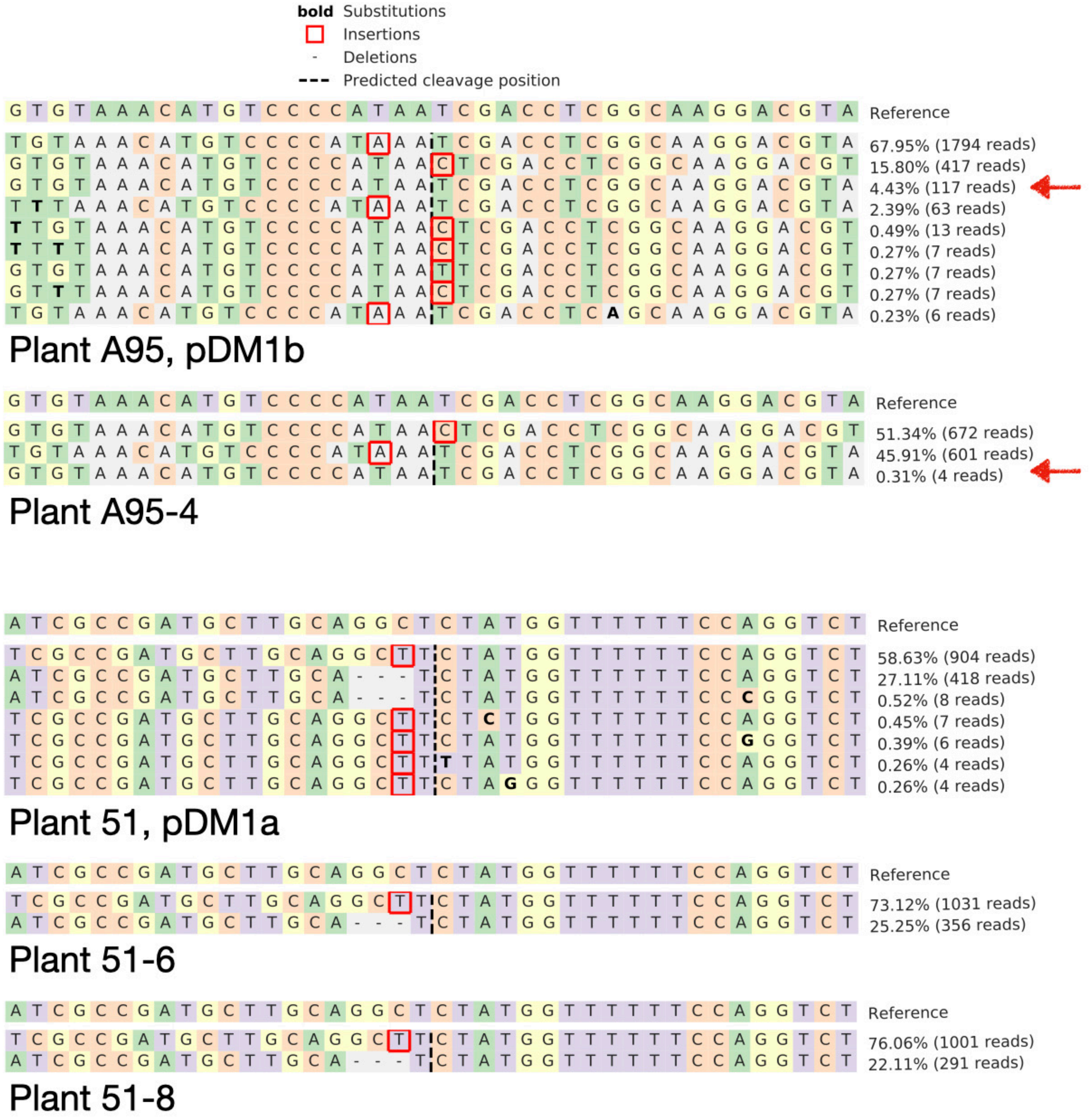
bold Substitutions
Insertions
Deletions
Predicted cleavage position
G T G T A A A C A T G T C C C C A T A A T C G A C C T C G G C A A G G A C G T A Reference
67.95% (1794 reads)
15.80% (417 reads)
4.43% (117 reads)
2.39% (63 reads)
0.49% (13 reads)
0.27% (7 reads)
0.27% (7 reads)
0.27% (7 reads)
0.23% (6 reads)
Plant A95, pDM1b
G T G T A A A C A T G T C C C C A T A A T C G A C C T C G G C A A G G A C G T A Reference
51.34% (672 reads)
45.91% (601 reads)
0.31% (4 reads)
Plant A95-4
A T C G C C G A T G C T T G C A G G C T C T A T G G T T T T T T C C A G G T C T Reference
58.63% (904 reads)
27.11% (418 reads)
0.52% (8 reads)
0.45% (7 reads)
0.39% (6 reads)
0.26% (4 reads)
0.26% (4 reads)
Plant 51, pDM1a
A T C G C C G A T G C T T G C A G G C T C T A T G G T T T T T T C C A G G T C T Reference
73.12% (1031 reads)
25.25% (356 reads)
Plant 51-6
A T C G C C G A T G C T T G C A G G C T C T A T G G T T T T T T C C A G G T C T Reference
76.06% (1001 reads)
22.11% (291 reads)
Plant 51-8

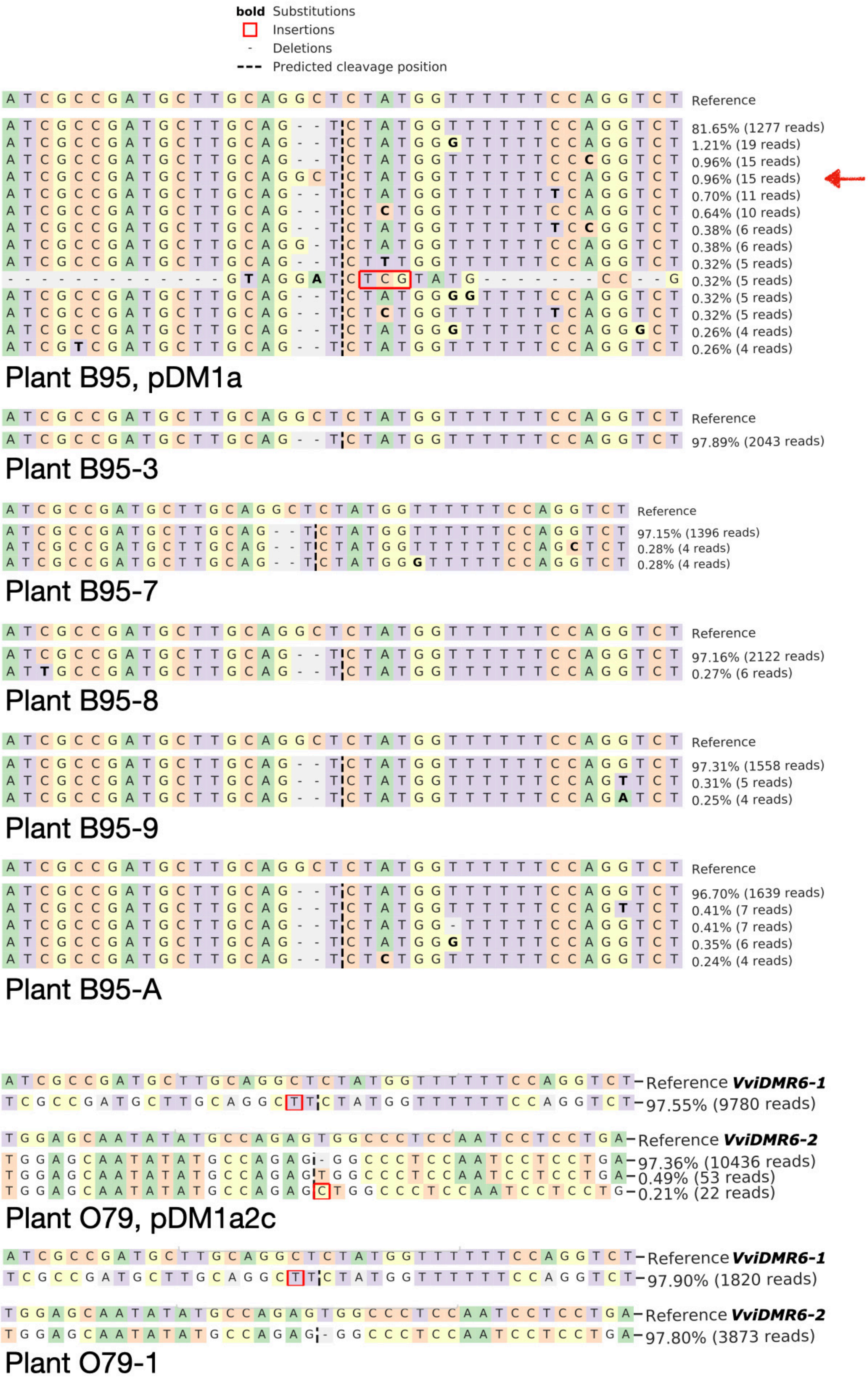
bold Substitutions
Insertions
Deletions
Predicted cleavage position
Reference
81.65% (1277 reads)
1.21% (19 reads)
0.96% (15 reads)
0.96% (15 reads)
0.70% (11 reads)
0.64% (10 reads)
0.38% (6 reads)
0.38% (6 reads)
0.32% (5 reads)
0.32% (5 reads)
0.32% (5 reads)
0.32% (5 reads)
0.26% (4 reads)
0.26% (4 reads)
Plant B95, pDM1a
Reference
97.89% (2043 reads)
Plant B95-3
Reference
97.15% (1396 reads)
0.28% (4 reads)
0.28% (4 reads)
Plant B95-7
Reference
97.16% (2122 reads)
0.27% (6 reads)
Plant B95-8
Reference
97.31% (1558 reads)
0.31% (5 reads)
0.25% (4 reads)
Plant B95-9
Reference
96.70% (1639 reads)
0.41% (7 reads)
0.41% (7 reads)
0.35% (6 reads)
0.24% (4 reads)
Plant B95-A
Reference VviDMR6-1
97.55% (9780 reads)
Reference VviDMR6-2
97.36% (10436 reads)
0.49% (53 reads)
0.21% (22 reads)
Plant O79, pDM1a2c
Reference VviDMR6-1
97.90% (1820 reads)
Reference VviDMR6-2
97.80% (3873 reads)
Plant O79-1
